# The emergence of dynamical instantaneous memory in the spontaneous activity of spatially confined neuronal assemblies *in vitro*

**DOI:** 10.1101/412320

**Authors:** Y. Piasetzky, M. Bisio, S. Kanner, M Goldin, M. Olivenbaum, E. Ben-Jacob, Y Hanein, M. Chiappalone, A. Barzilai, P. Bonifazi

## Abstract

Understanding the dynamics between communicating cell assemblies is essential for deciphering the neural code and identifying the mechanism underlying memory formation. In this work, in order to unveil possible emergent intrinsic memory phenomena in the communication between cell assemblies, we study the spontaneous dynamics of *in vitro* spatially confined inter-connected neuronal circuits grown on multi-electrode arrays. The spontaneous dynamics of the global network was characterized by the coupling of the activity independently generated by each circuit. The asymptotic functional connectivity of the network reflected its modular organization. Instantaneous functional connectivity maps on ten seconds epochs, revealed more complex dynamical states with the simultaneous activation of distinct circuits. When looking at the similarity of the generated network events, we observed that spontaneous network events occurring at temporal distances below two dozens of seconds had an average higher similarity compared to randomly played network events. Such a memory phenomenon was not observed in networks where spontaneous events were less frequent and in networks topologically organized as open lines. These results support the hypothesis that dynamical instantaneous memory, characterized by drifting network dynamics with decaying degree of similarity, is an intrinsic property of neuronal networks.

## Introduction

The nature of cortical representations is central for understanding how memory and cognitive operations are implemented by the brain. In 1949, Donald O. Hebb [1] introduced the concept of ‘cell assembly’ to describe a functional unit composed by a group of cells connected through excitatory synapses. Nowadays, thanks to more advanced techniques [2] evidences for the existence of Hebbian cell assemblies and the underlying associative learning mechanisms [3-5] have been provided. Indeed, multiple single-unit recordings revealed that the coordination of cell firing in cortical networks is more complex than postulated with an important role also played by the precise relative firing times and temporal correlations in neural ensembles [6-10]. All these observations led to a renewed wider vision of the neural coding problem, widening the rate coding to spike based coding mechanisms. Experimental and theoretical evidences support the possibility that the brain operates through coordinated activation of cell assemblies [11-15], which can be regarded as dynamic functional units that transiently interact with each other, shaping and underlying different brain states.

Despite the very recent technological progress [16-19], monitoring with high spatio-temporal resolution the structure and the dynamics of *in vivo* neuronal networks is still challenging. To this end, the development of *in vitro* neuronal networks is of significant interest due to their easy accessibility, monitoring, manipulation and modeling [20]. Moreover, in the last decades, many studies proposed advanced substrate patterning methods to support *in vitro* technologies, able to induce neuronal networks to grow according to an imposed topology: surface modification by silane chemistry [21], photolithographic techniques [22], deep-UV lithography [23], soft lithography [24] and spot-arrays of adhesion molecules [25, 26]

Thanks to those different patterning techniques, it has been possible to provide simplified but plausible representations of interacting cell assemblies through engineering inter-connected neuronal sub-populations [11, 12, 14, 27-30]. These experimental models constitute a valid alternative to widely studied 2D homogenous cultures which lack any spatial constraints on the self-organized emergent wiring of the circuits, a constraint that is definitely present in *in-vivo* architectures. Notably, using such *in vitro* structural modularity approach it has been shown how richer repertoires of spontaneous synchronizations emerge compared to 2D homogenous cultures [29-31].

Using such structured *in vitro* networks, it has been shown how different spontaneous motifs of activity can be found [32] and how memory can be imprinted by using focal chemical stimulations inducing local disinhibition on cultured neuronal networks. In [32], after an initial characterization of the network motifs and their initiation sites, the authors reported how a new motif starting at the site of chemical stimulation could be imprinted and played spontaneously without erasing preexisting motifs.

Similarity between spontaneous and evoked activity patterns was previously described in a number of *in vivo* studies [33] [34] and a recent study has confirmed such observation *in vitro* [35]. In this work, similarly to what previously described [12, 27, 29, 30] we employed polymeric structures made of polydimtheylsiloxane (PDMS) for confining neurons (Mata *et al.*, 2005; Liu *et al.*, 2005; del Campo & Greiner, 2007 Jackman *et al.*, 1999) to induce the self-organization of networks into inter-connected spatially defined neuronal assemblies. The developed modular networks have been cultured on conventional Micro Electrode Arrays in order to monitor their electrophysiological activity [36-39]. Focusing on the spontaneous occurrence of the network motifs in order to unveil possible emergent intrinsic memory phenomena in the communication between cell assemblies, we provide evidence about the existence of a short-term memory window lasting about two dozens of seconds within which motifs share high similarity. These results support the idea that instantaneous dynamical memory in brain networks represents an intrinsic capability of interconnected cell assemblies to maintain similar coordinated activation, and therefore holding the same information content, over time courses which have been typically reported in *in-vivo* working memory studies [40].

## Methods

### Ethics Statement

As experimental model for this research, primary cortical cultures from embryonic rats have been used in view of their large diffusion as animal model in neuroscience, in particular electrophysiology and MEA research. All procedures involving experimental animals were approved by the Italian Ministry of Health and Animal Care (authorization ID 023, April 15th, 2011).

### Primary cell culture

Cultures were prepared as described previously [12, 41]. Briefly, the entire neo-cortex of Sprague Dawley rat embryos (E18-19) was removed, chopped with scissors in a Trypsin EDTA solution. Dissociated cells were suspended in a growth medium (5% FCS, 2% B27, 1% L-Gln, 0.5% Pen-Strep in Neurobasal A medium). Cell suspension was diluted to reach the desired concentration.

### Neuronal network patterning using poly-d-lysine (PDL) stencils

The process of PDL patterning was detailed in previous publications [42] [29]. Briefly, PDL (Sigma, Cat. No. p7405) islands on top of MEAs were prepared with a soft lithography process using polydimethylsiloxane (PDMS) stencils. PDMS is an elastomer widely used for biomedical applications because of its well-known properties of highly temperature, chemical and oxidation resistance, biocompatibility, transparency and permeability to gases [43]; plus it is an electrical insulating and not toxic, making it suitable to the culture of primary neurons [44]. In addition, it can be easily micro-structured by soft lithography, thus obtaining several low-cost replicas from a single master [45-47]. An SU8-2075 (Micro Chem) mold was patterned on a silicon wafer. The pattern was identical to the negative pattern of the electrode array. The stencil was prepared by spin coating the wafer with PDMS. After detaching the PDMS substrate from the mold, the stencil was placed on commercial MEAs and the stencil’s pattern is aligned with the electrode locations. The PDL solution is dripped onto the PDMS stencil and incubated overnight at 37°C. The PDMS stencil is removed and MEAs were washed twice before cell plating. Once cells are plated, they spontaneously assemble to the coated islands and self-organize into active inter-connected circuits. For obtaining monolayers or clustered circuits as well as to impose and shape their connectivity, three parameters are required; cell plating density, distance between circuits, and circuits’ size. These parameters are not independent since, given a fixed distance between two circuits, the probability that they spontaneously establish a connection increase with the circuits’ size, by increasing the distance between the squares it was possible to change the probability to establish spontaneously generated inter-connections among the modules, passing from highly connected modules to isolated ones [42].

The cultures were maintained at 37°C with 5% CO2. The growth medium was partially replaced every 4 day. By increasing the distance between the squares it was possible to change the probability to establish spontaneously generated inter-connections among the modules, passing from highly connected modules to isolated ones.

### Electrical recordings

In this work, the commercial system purchased from Multi Channel Systems (MCS, Reutlingen, Germany) has been used. MCS provides different types of MEAs (in terms of electrode size and inter-electrode spacing), and the ones used here consist of 59 round electrodes made of TiN with three different layouts: *i) Standard MEA,* where electrodes are equidistantly positioned in an 8×8 layout grid, whose inter-electrode distance is 200 µm and microelectrode diameter 30 µm; *ii) 4QMEA1000,* which has a layout composed of four quadrants of microelectrodes with electrode spacing equal to 200 µm inside the quadrants and inter-quadrants distance of 1000 µm, with an additional central line of 7 microelectrodes (the electrodes diameter is 30 µm); and *iii) 60MEA500,* in which electrodes are equidistantly positioned in a 6×10 grid, whose inter-electrode spacing is 500 µm, and the electrode diameter 30 µm (Figure 1). For this study, we recorded and studied the activity of 21 networks between 21 DIV to 28 DIV. The number of neuronal assemblies per network were: 3 (N=3), 4 (N=10), 5 (N=6) and 6 (N=2).

**Figure 1.**
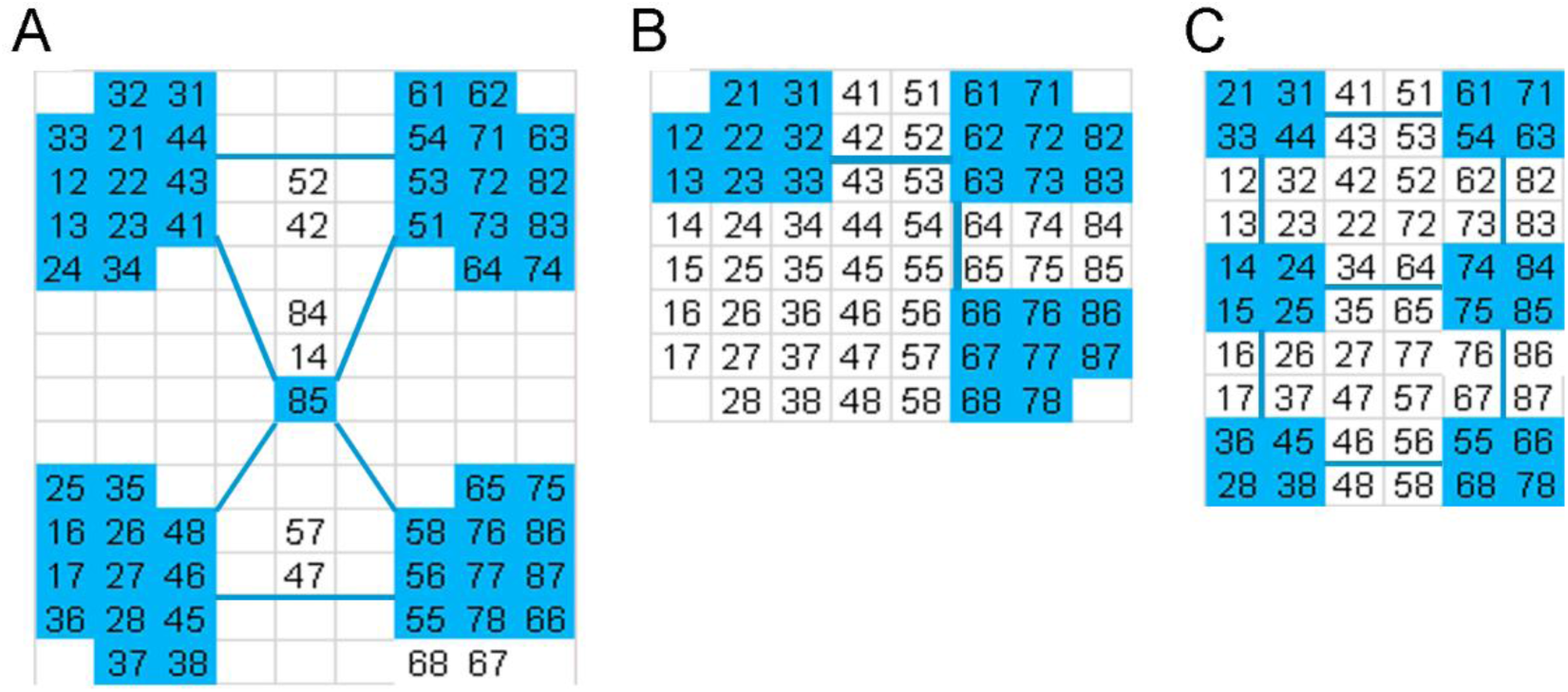
MEA layouts used and three representative network structures. (*A*) 60-4QMEA composed by four quadr electrode distance. 1 mm (with a 200 □m space between electrodes within the quadrants) and a central line of electrodes (with 200 and 300 □m distance). (B) Standard 8×8 MEA with 200 □m inter electrode distance. (C) 10×6 MEA with 500 □m inter-electrode distance. Blue areas marks recording electrodes beneath the cultured neuronal assemblies. Blue lines mark fiber bundles connecting distinct neuronal assemblies as visually detected by bright field microscopy. Self-organization of networks into inter-connected spatially defined neuronal assemblies was obtained using PDMS stencils (see Methods). Networks were composed by 6 neuronal assemblies or less.

### Experimental Protocol

The adopted experimental protocol was the following: *i)* 1-hour recording of spontaneous activity; *ii)* stimulation session, consisting of 5 minutes of stimulation (50 positive-then-negative pulses are delivered at 0.2 Hz, amplitude 1.5 Vpp, duration 500 µs and duty cycle 50%) delivered to each module (i.e. two channels per module) followed by 10 minutes of rest. For each assembly, a random electrode producing clear within-module evoked activity, was chosen for studying the evoked network response. The purpose of the stimulation was not to imprint memory, rather to evaluate the effective connectivity (i.e. propagation of activity) between the circuits.

### Data analysis

All data analysis has been performed using MATLAB (MathWorks, Natick, MA).

#### High-pass filtering, spike extraction, active electrodes (AEs) and multi-unit activity (MUA)

Electrical recordings were first pre-processed using a zero-phase digital filtering (“filtfilt” function) with high-pass cut-off set at 100 Hz. All analysis steps described below were applied to the filtered signals. For each electrode, the noise level was estimated by fitting the probability density function of the signal with a Gaussian, in the 5^th^ to 95^th^ percentile interval, over the initial ten minutes’ recordings. A threshold of 4 standard deviations below the mean was used to extract the timing of the negative peak of the spikes with an imposed refractory time between spikes of 1 ms. In this work, no spike sorting was applied and all the spikes, i.e. multi-unit activity (MUA), recorded by each electrode were considered. In particular, only electrodes recording spikes with a total average frequency higher than 0.001 Hz were included in the analysis, and they are referred in the text as active electrodes (AEs).

#### Instantaneous firing rate and functional connectivity

The instantaneous firing rate vectors (*IFR*(*t*)) representing the activity recorded by the AEs in a giving time frame *t*, was calculated in time bins of 250 ms, and each scalar of the vector (*IFRn*(*t*), with 1<=n<=N where N is the total number of AEs) represented the number of spikes detected in the corresponding electrode. The asymptotic functional connectivity (a-FC) matrix was calculated electrode-wise as the Pearson correlation between IFR time series of the electrodes. Specifically, given a pair of electrodes (i,j), the correlation between *IFRi*(*t*) and *IFRj*(*t*) was computed. The same procedure was applied for computing the instantaneous functional connectivity matrix (i-FC), but subdividing the time series into consecutive windows of ten seconds duration.

The IFR similarity matrix was calculated as the cosine between the IFR vectors. For a given set of T time frames, the IFR matrix had a size of TxT. The k-order diagonal (with k positive integer) of the IFR matrix was defined as the set of elements (j,j+k) with j>=1 and j<=T-k.

#### Bimodal lognormal distribution of inter-spike interval and inter-spike interval cut-off threshold (COT)

For each network, the global inter-spike interval sequence was calculated by pooling all the inter-spike intervals of each active electrode. The distribution of the decimal logarithm of the global inter-spike intervals (in ms), calculated in bins of 0.05 in the interval 0 (corresponding to 1 ms) to 5 (corresponding to 10000 ms), was fitted by the sum of two lognormal distributions. The time corresponding to the minimum in between the two lognormal was applied as cut-off threshold (*COT*) to remove “isolated” spikes, i.e. preceded or followed by spikes with an interval larger than *COT*.

#### Detection of network events (NEs) and conversion into activation-order matrixes (AOM)

In order to detect NEs, a binary time-series with a time resolution of one millisecond was built for each electrode. In order to mark time windows with neuronal firing bursts, epochs of inter-spike intervals below *COT* were marked with ones. A NE was considered to start when more than one electrode was firing at the same time and stopped when less than two electrodes were simultaneously firing. Each NEm (*1<=m<=M*, where M is the total number of NEs) was represented by an activation order matrix *(AOM*) of size NxN, where N are the number of AEs in the experimental recording. The element *AOM(m)_ij_* was set to 1 (−1) if, in the NE_m_, the firing of electrode *i (j)* preceded the one of electrode *j (i)*. If either *i* or *j* did not participate in NE_m_, the elements *AOM(m)_ij_* and *AOM(m)_ji_* were set to zero.

#### NEs similarity and average similarity of NEs as function of temporal distance (SNE)

For every pair *(i,j)* of NEs, the similarity matrix *S_ij_ (1<=i,j<=M*, where M is the total number of NEs) was calculated according to the following:

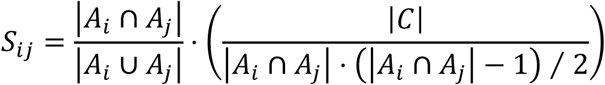

where A_i_ is the set of electrodes that participate in event *i*, A_j_ is the set of electrodes that participate in event *j, C* is the number of identical non-zero elements in AOM(i) and AOM(j), | · | denotes the cardinality of a set, and B is the cardinality of the intersection of A_i_ and A_j_ (i.e. B=| A_i_ ∩ A_j_ |). This definition of similarity which varies between 0 (no similarity) and 1 (identical NEs), quantifies in a unique parameter the similarity of the patterns of electrodes recruited in the NEs (spatial component) and the amount of identical pair-wise activations (temporal components). When dendrogram analysis on the similarity matrix was performed, Euclidean distance and Ward’s linkage criteria were applied.

In order to calculate the average similarity between pair *(i,j)* of NEs as a function of their temporal distance Δt (SNE(Δt), where Δt=|ti-tj|, and t_i_ and t_j_ are the time in seconds of NE_i_ and NE_j_), the time window [0 ÷ 50sec] was binned in one second intervals and the average similarity of every pair of NEs with Δt falling within a given bin was calculated.

In order to calculate the SNE on randomized sequence of NEs, the order of the NEs was randomized while keeping the AOM and time of NE occurrence t_i_ (1<=i<=M) unchanged. The average and standard deviation of the mean.

#### Assembly-pooled PSTHs and latency of response

Assembly-pooled PSTHs (Post Stimulus Time Histogram) with ten milliseconds bins were obtained by pooling all the spikes recorded within a given assembly in response to electrical stimuli. The latency of the PSTH peak was evaluated in sliding windows for obtaining higher temporal accuracy. Z-score was used to asses which neuronal assembly responded reliably to the stimulation.

## Results

### Spatially confined inter-connected neuronal circuitries: spontaneous dynamics and structural – functional match

We used *in vitro* cultured neuronal circuitries with predefined architectures (see Methods).

In this work, we used networks with different architectures grown on MEAs with the electrode layout optimizing the interface of the networks (Figure 1). In all the experiments described in this work, the spontaneous activity was first recorded one hour, followed by a stimulation session where at least one of the electrodes recording from each module was used to evoke intra-circuit activity and to test the effective connectivity (i.e. propagation of activity) between the circuits (see Methods and Figure 2). All engineered neuronal networks used in this work behave as weakly coupled neuronal circuitries capable of independently sustaining synchronizations and occasionally synchronizing between them (Figure 3).

**Figure 2.**
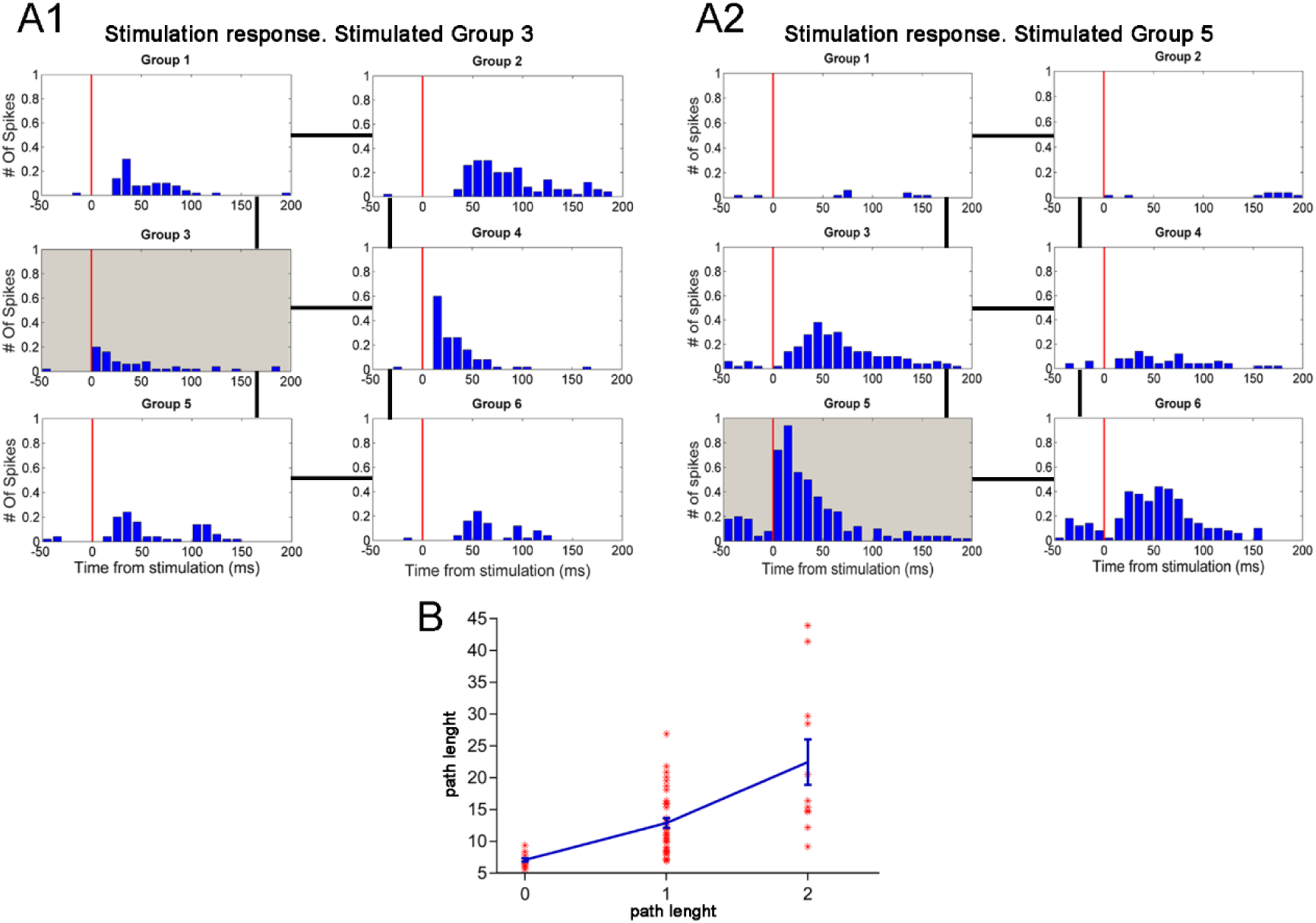
Assembly-pooled PSTHs for probing effective connectivity. *(A1-A2)* Assembly-pooled PSTHs (with ten milliseconds bins) were obtained by pooling all the spikes recorded within a given assembly in response to electrical stimuli. A sequence of fifty stimuli (see Methods) was applied to each neuronal assembly and the response of the different assemblies was measured by PSTHs. For each assembly, a random electrode producing clear within-module evoked activity, was chosen for studying the evoked network response. In the panels *A1* and *A2*, the PSTHs are organized as the network topology, where lines mark the connections between assemblies. Vertical red lines mark the time of stimulation. Gray PSTHs mark the module where the stimulaiton was applied. Note that modules directly connected to the stimulated one respond more reliably and with an average shorter time latency. (*B*) The latency of the PSTH peak (evaluated in sliding windows for obtaining higher accuracy) is plotted against the path length, i.e. the number of links separating pairs of assemblies. The vertical bars represent mean ± SEM.

**Figure 3.**
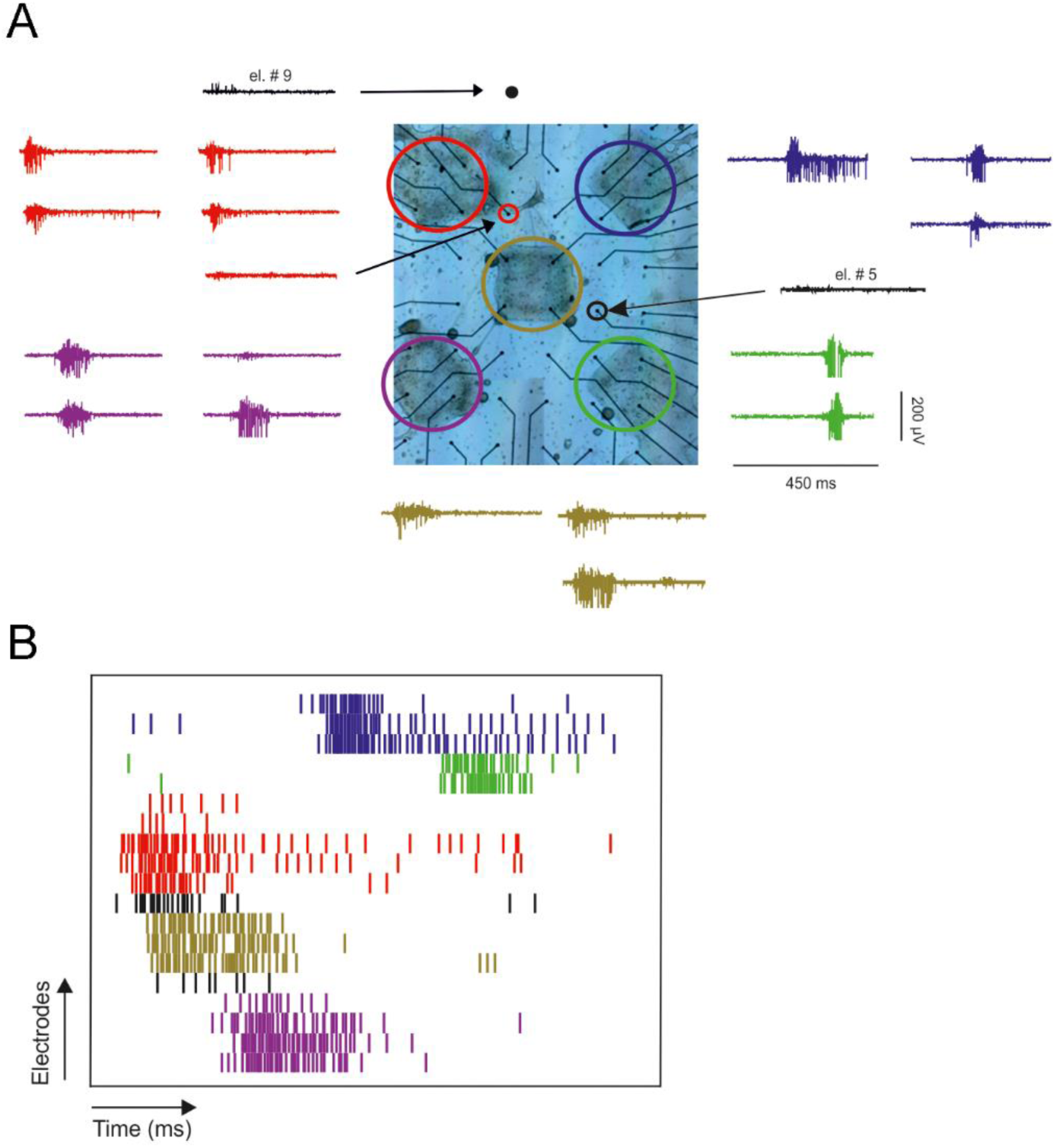
Structure and dynamics of a representative patterned network. (*A*) Bright field image of the network grown on 6×10 multi-electrode array with 500 inter-electrode distance. The large colored circles highlight the spatially confined interconnect, self-organized neuronal circuits. Electrical traces of 500 ms duration recorded from the different electrodes located under the assemblies are shown with the same color code. Black arrows links traces to the few electrodes that were located out of the circuits but still capturing neuronal signals. (*B*) Raster plot of the activity shown in panel A. Note the network activity pattern is composed by synchronous firing in each neuronal assembly.

In order to characterize the spontaneous synchronizations, we first calculated the instantaneous electrode firing rate (IFR_n_(t), where 1<=n<=N, N is the number of active electrodes and *t* is the time frame of the recording; active electrodes abbreviated as AEs, are electrodes capturing the activity of the neurons recorded in different location of the networks, through the MEA. In particular, for each electrode we considered the multi-unit activity (MUA) possibly originating from a few neurons, and we counted the number of spikes in time windows of 250 ms, i.e. the characteristic temporal duration of network synchronizations [48]. Next, we reconstructed the functional connectivity (FC) of the network, i.e. the statistical dependency of the activity fluctuations, by computing the Pearson correlation between the IFR_n_(t) for every possible pair of active electrodes. When the correlation was calculated over the entire session of spontaneous recording (about one hour), the FC organization fully reflected the structural organization of the network (Figure 4*A*), with a significant difference (Kolmogorov-Smirnov test, p<0.01) between the average intra-circuit FC of 0.35 +/- 0.08 and inter-circuit FC of 0.22 +/- 0.05 (N=21, standard deviation of the mean; for DIV and neueonal assemblies per network see Methods). We can consider this as an asymptotic or static representation of the FC of the network (a-FC), where the modular organization of the a-FC matrix reflects the modular organization of the neuronal network, with higher structural and functional connectivity within each circuitry, and lower connectivity between them. Note that the order of the electrodes in the FC matrix of Figure 4*A* has been appositely arranged so that electrodes recording from the same circuit are close to each other in the electrode sequence.

**Figure 4.**
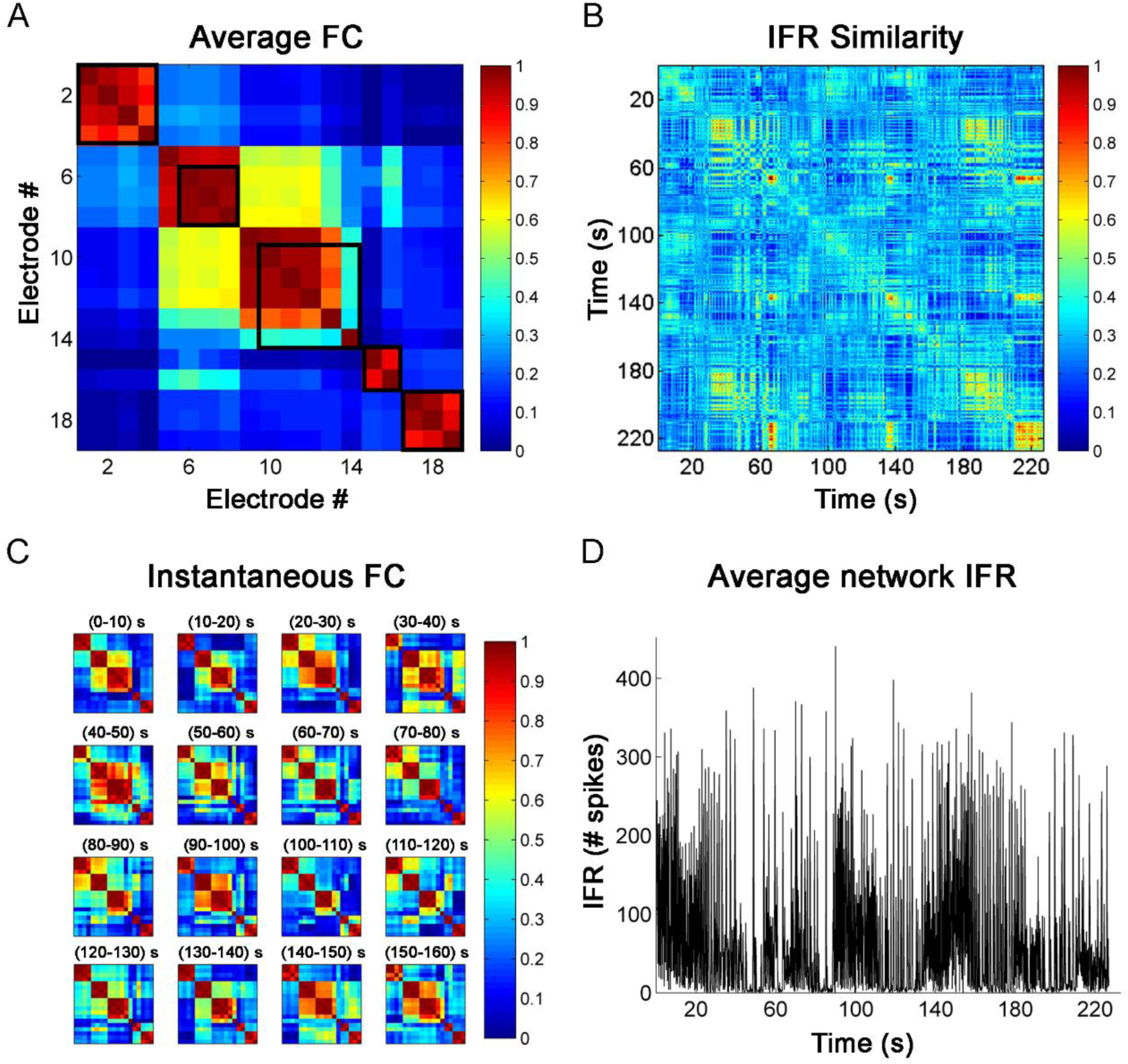
Asymptotic and instantaneous functional organization. (*A*) Asymptotic functional connectivity matrix between the instantaneous firing rate recorded by the electrodes. The IFR was estimated in time bins of 250 ms and the Pearson correlation between the IFR time series was calculated over an entire session of spontaneous activity lasting about an hour. The network is the same shown in Figure 3 and the black squares mark the electrode beneath the same neuronal assembly, with the same labeling used in Figure 3. (*B*) Similarity matrix between the IFR vectors (where each element represents a recording electrode) for a representative period of about 220 s. The cosine similarity was used (see Methods). (*C*) Same as panel A but the functional connectivity has been calculated instantaneously in time windows of 10 seconds. Note both the different dynamical states, composed by functionally connected modules, explored by the network and the consistent high functional connectivity within each module. (*D*) Average network IFR, i.e. obtained by summing the elements of the IFR vector in each time bin, for the time window reported in B.

When instantaneously computing the FC on shorter time epochs, i.e. in the order of seconds, which is the characteristic temporal scale of the inter-burst events [48], the instantaneous FC matrix (i-FC) displayed very different characteristics (Figure 4*B*). The i-FC shows that the network spontaneously explores different states composed by the correlated activation of distinct electrodes and circuits. Specifically, when looking at the inter-circuit FC, the ratio between i-FC and a-FC was in average 1.5+/-0.2 (N=21).

When looking at the similarity of the IFR_n_(t) at different time frames, we observed the presence of similar profile of network activity in closer time frames (i.e. next to the diagonal of the similarity matrix shown in Figure 4*B*, see Figure 5) but also at much larger time windows. Figure 4*D* reports the average network IFR for the different time frames considered in Figure 4*B*. The average along different diagonals shows an initial higher similarity transient in the first two dozens of seconds before decaying to baseline levels (Figure 5). This observation is the rationale of the analysis reported below, aimed at dissecting the dynamics in the similarity of the spontaneous events as a function of their temporal distance, prompting the existence of a temporal window lasting dozens of seconds where the network shows a memory of the previously displayed events.

**Figure 5.**
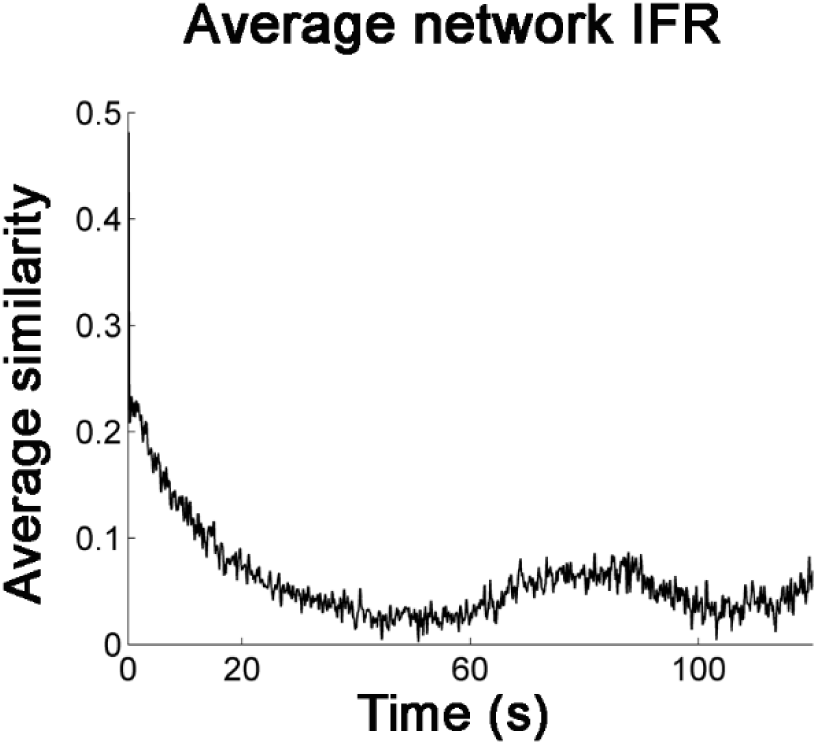
Average similarity between IFR vectors separated by a given time interval. Given the IFR similarity matrix of size SxS (as shown in Figure 4*B*), the *k*-order diagonal (with k positive integer) was defined as the set of elements (j,j+k) with j>=1 and j<=S-k. The average of the elements in the *k*-order diagonal represents the average similarity of all IFR vectors separated by a fixed time interval of k*250ms (since the IFR is calculated in 250ms time bins, see Methods). The average of the diagonals of the matrix shown in Figure 4*B* is here plotted.

### Network events: identification and similarity

In order to get deeper into the instantaneous spatial – temporal organization of the network activity and its dynamics over the entire recording, we first focused on the temporal features of the MUA (Figure 6*A*). When looking at the inter-spike-interval over the entire spontaneous activity recording pooled across all electrodes, we observed that it is distributed as a bimodal lognormal distribution (Figure 6*A*), which highlight the existence of two characteristic temporal scales, a fast scale in the order of 10 ms (100 Hz fast firing) occurring during network synchronizations [29] and a slow temporal scale in the order of 1 sec (1 Hz slow firing) reflecting the interval between network synchronizations. As previously reported in literature, the fast temporal scale dynamic corresponds to network bursts or neuronal synchronizations while the slow one characterizes the inter-burst epochs [49]. In order to identify network events (NEs)composed by neuronal bursts and synchronizations recruiting more than one circuit, we first discarded out-of-burst spikes (see Methods) by applying the cut-off-threshold (*COT*) identified as the minimum between the two lognormal distributions (Figure 6), on the inter-spike-intervals. Once out-of-burst spikes were discarded, neuronal bursts epochs were identified in each electrode (Figure 6*B*; see Methods) by binarizing the time windows during which the inter-spike-interval was lower than the *COT*. A time resolution of one millisecond was used in the binary electrode time series where ones marked the bursting windows. According to the definition used in this work, a NE started when more than one electrode was bursting and stopped when less than two electrodes were bursting simultaneously. Every NE was represented by a NxN activation-order matrix (AOM, Figure 6*C*), where N was the number of AEs in the experiment and where every element (i,j) (with i,j<=N) reported the pair-wise order of activation in the NE (one if i anticipated j, minus one if j anticipated i and zero if either i or j did not participate in the NE; AOM is antisymmetric by definition).

**Figure 6.**
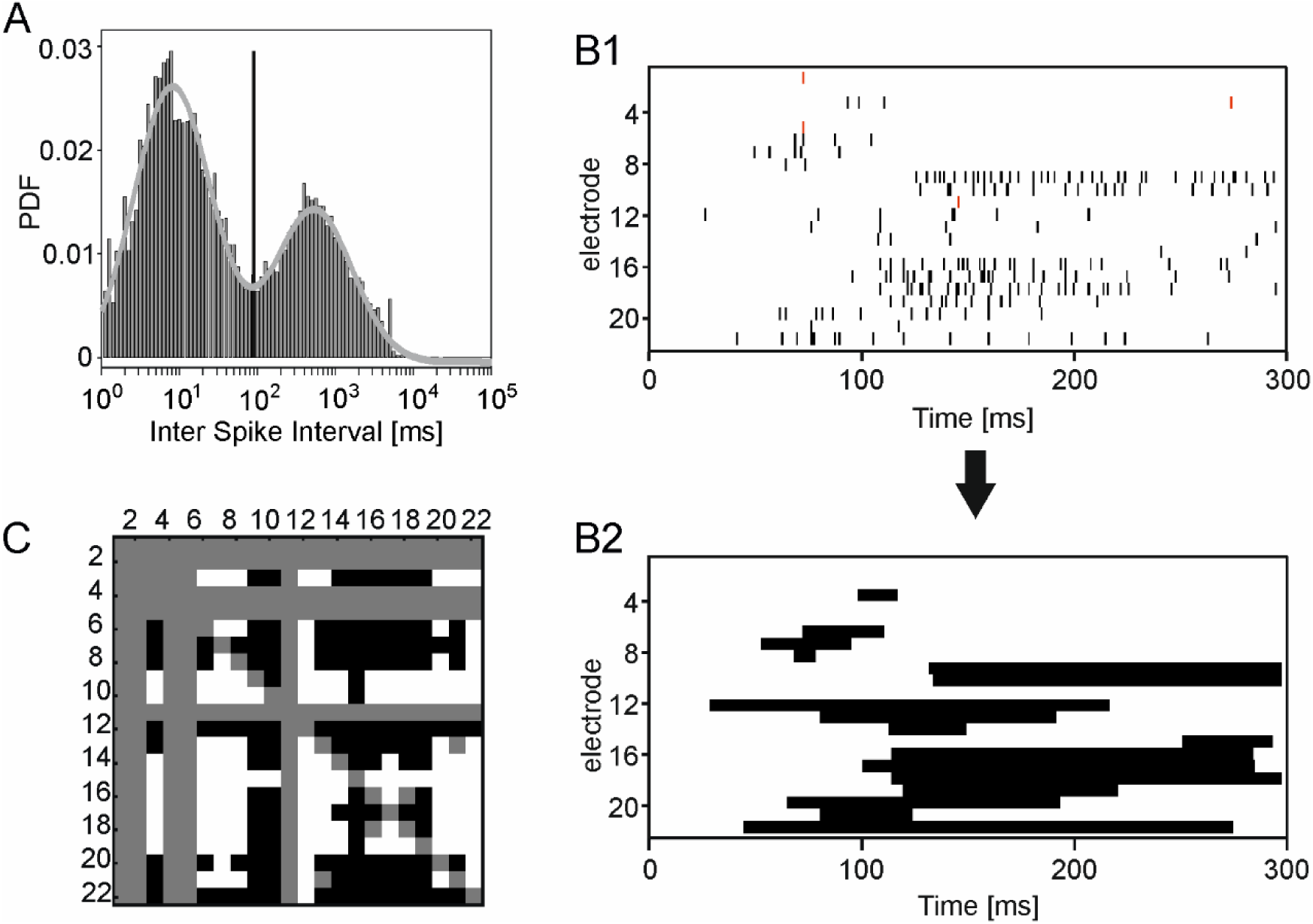
Representing network events as activation-matrix order. (*A*) Pooled inter-spike distribution for a representative network (the same show in Figure 3). All inter-spike intervals recorded by all electrode over an entire session of spontaneous activity have been pooled into the statistic. Note the use of the logarithmic time scale (x-axis). The fit with the sum of two lognormal distributions is shown with the gray line. The vertical black line highlight the time of the minimum between the lognormal fit, which has been used low pass cut-off. (*B1*) Spike trains of a representative network events. Red lines mark the spikes discarded by the cutoff procedure. (*B2*) Binarization of the spiking activity for the network event reported in B1. Period of firing bursts (i.e. with an inter-spike interval below the threshold marked in panel A) are marked as ones (black horizontal bars). (*C*) The activation order of the network event shown in the panels B is represented by the activation-matrix order. Gray rows and columns correspond to non-firing electrodes. A black (white) square for the element (i,j) corresponding to the firing of electrode i anticipating (following) the firing of electrode j based on the binarized activity of panel B2.

Given the above NE temporal representations we calculated the pair-wise similarity between all NEs (Figure 7). The similarity index used in this work (see methods) takes into account the similarity of the spatial components of the NEs (given by the Jaccard index, i.e. the size of the intersection over the union of the sets of electrodes recruited in the NE pair) and the similarity of the temporal components of the NEs (i.e. the normalized number of matching elements in the AOMs). Figure 7 shows the similarity matrix for a representative network, where the NE indexes are reordered after clustering similar NEs (using dendrogram analysis, see methods). The clustering reveals the existence of groups of similar NEs (squared structures along the diagonal of the matrix), i.e. NEs with both high spatial and temporal similarity spontaneously appear.

**Figure 7.**
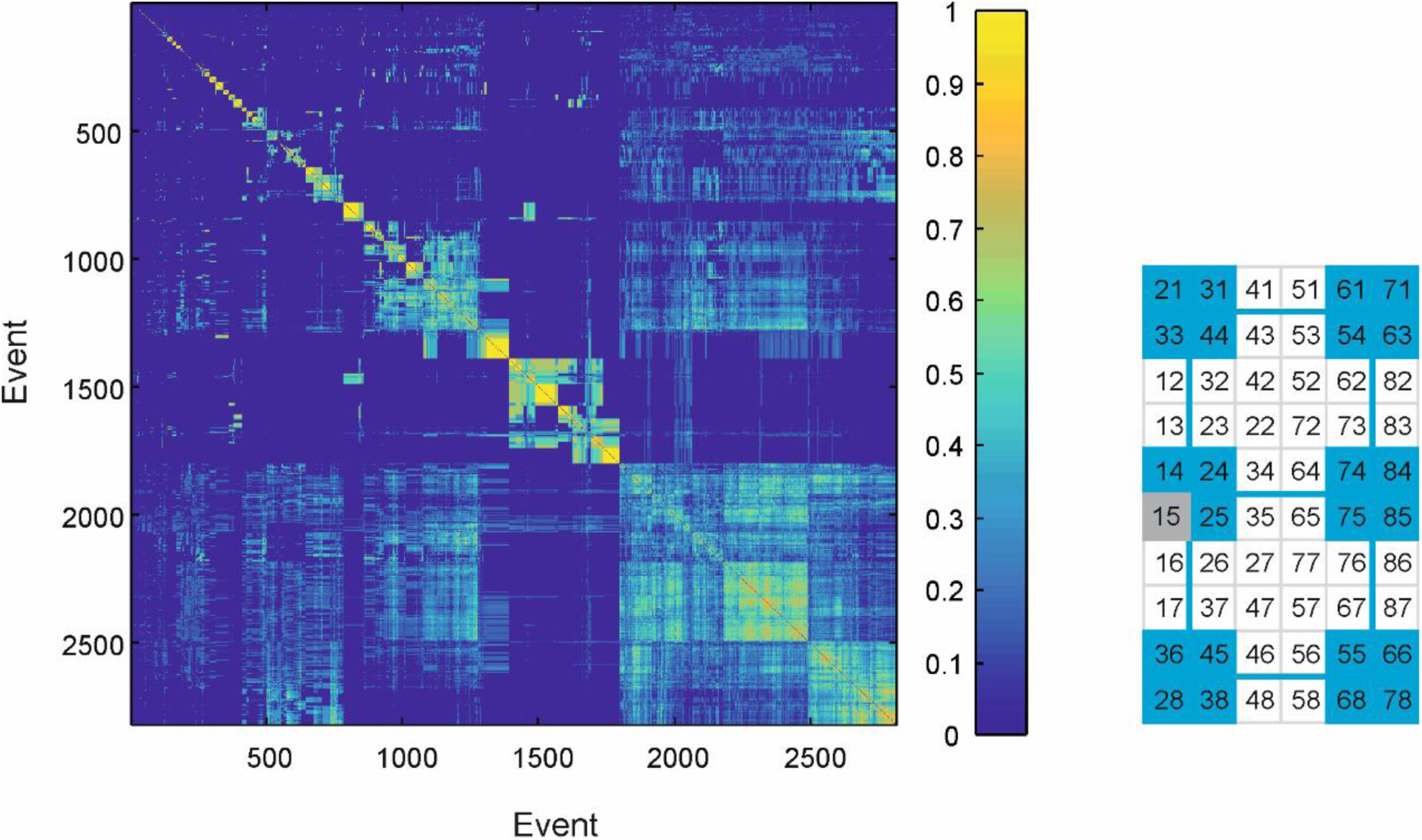
Similarity between network events. Similarity between network events (see Methods) recorded in the schematized network shown on the right side. Groups of similar events appear as squares in the diagonal of the matrix.

Finally, we investigated how NEs similarity evolves in time. Therefore, we calculated the average NEs similarity as a function of their temporal distance *(SNE(Δt))*. The results of this analysis revealed the existence (in 15 out of a total 21 networks) of a time window of 22.7+/-3.4 seconds within which the similarity of the NEs is higher than what expected by randomly reordering NEs. This memory-like phenomenon occurred only in networks whose structural topology was not chain-like, i.e. organized as an open one-dimensional line (5 out of 21 studied networks) (Figure 8). In addition, this phenomenon did not occur on one network with an average inter-NE interval larger than 30 seconds.

**Figure 8.**
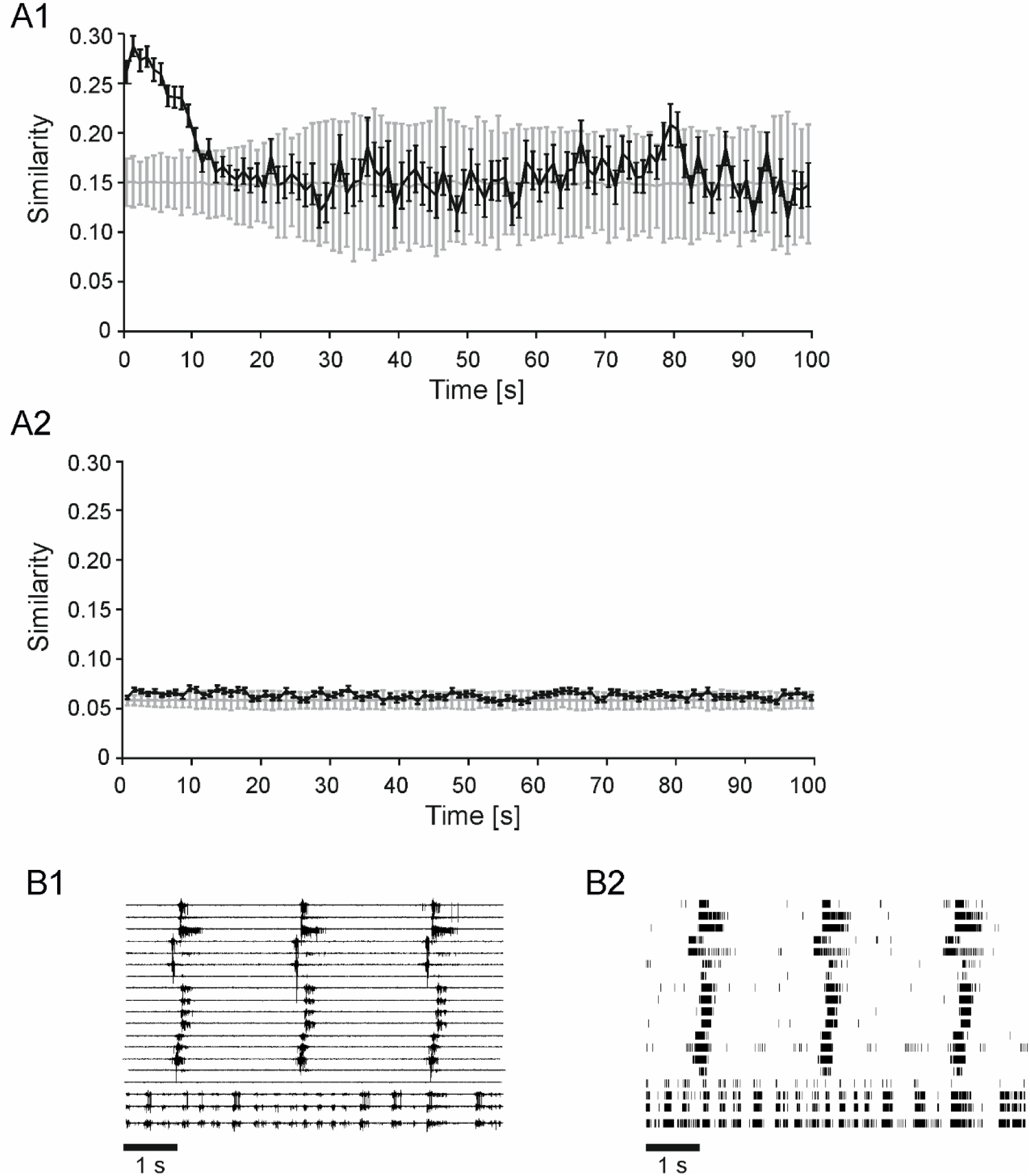
High similarity between close-in-time spontaneous events in non-chain networks. (*A1*) Emergence of instantaneous dynamical memory for a representative non-chain network. In black is shown the average similarity between events as a function of the inter-event distance. In grey the same analysis performed on randomly reshuffled events. (*A2*) The same as panel A1 but for a network organized as an open chain. (*B*) Electrical traces ad raster plots of three similar events played consecutively by the network shown in A1. Note the time delay of about 2.5 seconds between them. In these events, only four out of five neuronal assemblies participated with the same activation order. The bottom traces correspond to the non-participating assembly.

## DISCUSSION

In this paper, interconnected neuronal sub-populations have been used in order to investigate how their spontaneous dynamics (i.e. motifs of activity) relates to their particular topology, and to test the hypothesis of a memory mechanism in which similar spatiotemporal dynamical patterns are observed.

Although very informative, *in vivo* experiments do not allow a controlled manipulation of the spatio-temporal dynamics of neuronal networks. On the contrary, i*n vitro* systems, constitute a successful experimental model of neuronal dynamics (Orlandi Nature Physics 2013), and can be easily accessed, manipulated and monitored (Bonifazi et al., 2013). Therefore, in order to provide a simplified but plausible representation of the intrinsic modularity of the nervous system, we realized a multi-modular system, thus reproducing *in vitro* a group of ‘neuronal assemblies’, defined, similarly to Hebb’s view [1], as a set of anatomically and functionally connected neurons [5].

Several methodologies have been recently presented in the literature to drive the growth of neuronal cells according to pre-defined topologies. More specifically, substrate patterning methods were introduced with different aims: *i)* for selective positioning of cells through, e.g. chemical and mechanical guiding of their extensions [50] [51] and the use of physical constraints [52]; *ii)* for selective control of neurites guidance through, e.g. un-modified collagen scaffolds [53] or by controlling the direction of signal propagation at single-cell resolution [54]; and *iii)* for studying the functional properties of networks with imposed topologies [11, 55, 56] by inducing neuronal networks to develop a range of predefined modular structures [14].

Indeed, the networks realized in the context of this work can be defined as ‘multi-modular’ and were composed of spatially confined neuronal circuits. The rationale of this approach originates from the following argument. Uniform, i.e. spatially unconstrained, neuronal cultures generate stereotyped synchronized events recruiting the all neuronal population [48, 57]. Such spontaneous synchronizations are also present when the network scales down to a population size of few dozens of cells [27, 30]. Therefore, when *in vitro* self-organized networks of different population sizes do not experience any connectivity constraints, being axons and dendrites capable to grow freely also over long-distances, they naturally display stereotyped globally synchronized dynamics [48, 57]. Indeed, in the culturing two dimensional space, there is not any a-priori imposed economy principle of connectivity between cells, and this represents a very different condition compared to in-vivo where spatial constraints affect networks’ topology. By imposing spatial constraints on the two-dimensional space, spatially distinct circuits are created, and can independently generate spontaneous synchronizations while their mutual synchronization is strictly dependent on their connectivity strength. Specifically, poorly connected circuits occasionally synchronize while strongly connected circuits generate synchronized events recruiting the entire neuronal population as observed in uniform networks [58]. Therefore, the patterning design was mostly affected by two parameters, the circuit area and the inter-circuit distance, whose combination allowed to maintain the mutual synchronization between the spatially defined neuronal circuitries while preserving their independent local spontaneous dynamics [42].

Different structural layouts composing multi-modular networks (e.g. chain-like and feedforward networks) have been experimentally used to investigate how the layout influences the spontaneous activity. Furthermore, these networks have the capability of self-organizing forming spontaneous connections among the modules with the possibility to study how dynamics changes compared to uniform networks and to networks composed of well-known anatomical layouts [59]. Multi-modular networks can be electrically stimulated with the possibility to have a better control over the evoked activity, thanks to the structural and functional confinement ensured by the neuronal circuits. Several studies have been conducted within this framework, in particular, for investigating the relationship between structural and functional connectivity [60], and for understanding the role played by each circuit in processing neural information [61] [62].

Therefore, the multi-modular modular neuronal networks adopted in this work provided a powerful tool for investigating the dynamics of communications among neuronal assemblies and the capability of structurally defined neuronal networks to generate large repertoire of dynamical motifs.

When looking at the similarity of the spontaneously generated NEs over time, we observed that network events displayed higher similarity within a time window of about twenty seconds, compared to randomly played (i.e. reshuffled) network events. This instantaneous memory-like phenomena shares similarity with the working memory (WM) process studied *in-vivo* in intact neuronal circuitries [40]. Although the dynamics described in this work are spontaneous and not elicited by stimulations, the endogenous instantaneous memory and the WM mechanisms share important similarities; Network models trying to reproduce WM phenomena have shown that network activity behaves as a continuous attractor, where the bump of the population firing rate survives the disappearance of the stimulus [65], in a sort of sustained spontaneous activity shaped by the experienced stimulation and ii) a localized persistent activity pattern elicited by the stimulation tends to drift randomly as a diffusion process during the delay period [66], coding for a distinct but similar stimulus (specifically with slightly drifted spatial location).

The drift of the population activity bump observed in WM models [65] is conceptually similar to the decay of similarity between network events reported in this work. In analogy to the instantaneous memory phenomenon reported in our research, the variability of discharges from single neurons and the level of correlation between neurons recorded simultaneously during the WM tasks, vary consistently with the drifting bump model [65] and contrary to the idea that inaccuracy in WM is primarily due to the slow decaying of the bump of neuronal activity itself.

The fact that spontaneous dynamics can be a grounding mechanism for working memory is furtherly supported by the evidence that similarity between spontaneous and evoked stimulated activity patterns has been reported in a variety of *in vivo* [33, 34, 68-70] and *in vitro* [71] studies.

This work describes memory emerges in a non-chain like networks, this matter is interpretable by the need of recurrent connectivity allowing the reverberation of activity.

We propose that the spontaneous activity of a network reflect the current state of the network which slightly change and naturally drift over a time course of two dozens of seconds, and suggest an interesting future approach that the state of the network, which shape the spontaneous activity, might be shaped by external stimulations (environment).

## Acknowledgments

P.B. received EU funding from ICT-FET FP7 (Young Explorers; Grant No. 284772). P.B. also acknowledges financial support from the Ikerbasque Foundation and from the MINECO (Spain, grant SAF2015-69484-R). A.B. received funding from GIF (I-192-418) and ISF (421/15).

